# Illuminating Glucomannan Synthases to Explore Cell Wall Synthesis Bottlenecks

**DOI:** 10.1101/2025.03.09.642175

**Authors:** Annika Grieß-Osowski, Madalen Robert, Moni Qiande, Stefanie Clauss, Cătălin Voiniciuc

## Abstract

Hemicelluloses are important dietary fibers and a key component of lignocellulosic biomass. Despite numerous observations for fluorescently tagged cellulose synthases, the subcellular journeys and biochemical activities of intracellular cellulose synthase-like enzymes such as β-mannan synthases (ManS) remain largely unexplored. This study identifies C-terminal fluorescent protein tags that maintain ManS activity in the yeast to accelerate the Design, Build, Test, Learn cycles for polysaccharide biosynthesis. Using the *Amorphophallus konjac* ManS as a case study, we demonstrate that the enzyme co-localizes with a known yeast marker for the Golgi apparatus despite the toxic effects of plant glucomannan accumulation in *Pichia pastoris*. The ManS first transmembrane domain was found to be critical for the punctate localization of the enzyme, its overall expression level and its function. Additionally, we explored how fluorescently tagged ManS is influenced by genetic or chemical perturbations of native yeast cell wall components, such as reducing protein mannosylation and severely disrupting β-1,3-glucans. Finally, we identified alternative feeding strategies and episomal vectors for *Pichia*, which were extended to *Saccharomyces cerevisiae*, to accelerate hemicellulose research. We propose that expanding the Plant MoClo-compatible plasmid repertoire is essential to swiftly prototype carbohydrate-active enzymes in yeast before proceeding with more time-intensive analyses in plants. Requiring only hours or days instead of weeks or months for plant transformation/regeneration, our yeast prototyping strategies can de-risk the bioengineering of carbohydrate-active enzymes.

## INTRODUCTION

Enzymes from the cellulose synthase (CESA) superfamily elongate cellulose and most hemicelluloses,^1^ which collectively form the main network of fibers in plant cell walls. In the last two decades, N-terminal fluorescent protein (FP) tags on CESA proteins became popular tools to study cell wall dynamics and have revealed molecular insights into the localization and trafficking of CESA complexes *in planta.*^2–4^ While FP-CESAs are only active once secreted to the plasma membrane, CESA-like (CSL) enzymes producing β-1,4-linked mannans (CSLA), xyloglucan (CSLC), or mixed-linkage glucans (CSLF, CSLH, and CSLJ) function predominantly in the Golgi apparatus and their products are delivered extracellularly.^5^ However, the subcellular journeys and biochemical activities of most CSL enzymes have remained hidden, limiting long-standing aims to increase hexose sugar content (e.g. glucose, Glc, and mannose, Man) in bioenergy crops. For example, a prior attempt to boost mannan production in seeds through CSLA overexpression strategies not only failed to enhance cell wall composition,^6^ but caused severe genetic and metabolic perturbations.

Since plant transformation remains slow compared to microbial bioengineering, surrogate hosts are needed to accelerate Design, Built, Test, Learn cycles for polysaccharide biosynthesis or modification.^7^ Recently, *Pichia pastoris* (also known as *Komagataella phaffii*) has emerged as a promising yeast cell factory to study how plant CSLAs produce (gluco)mannans.^8^ While classical *in vitro* assays require microsomal extracts and radiolabeled nucleotide sugar donors,^9,10^ endogenous precursors in *Pichia* cells are sufficient for plant β-1,4-mannan and β-1,4-glucan synthases and their co-factors.^8,11,12^ Nevertheless, the attachment of superfolder GFP (sfGFP) to the N-terminus of β-mannan synthases such as the *Amorphophallus konjac* AkCSLA3, referred to as ManS for the remainder of the article, significantly reduced or abolished activity in yeast. Here, we explore the influence of FPs at the C-terminal end of ManS and extend the portfolio of genetic and chemical tools available to investigate plant hemicellulose production in yeast cells.

## RESULTS AND DISCUSSION

### Building A Set of Functional ManS-FP Fusions

First, we generated a ManS-sfGFP containing C-terminal tag in the pPICZ B vector, and compared it to the previously tested^8^ sfGFP-ManS. Seeking to determine the importance of induction time, cells were first grown in glycerol-containing medium for biomass accumulation. The sfGFP control strain and new ManS-sfGFP lines showed significant expression after 6 h of methanol induction (and peaked within 9 h) based on FP quantification using a plate reader (**Figure 1A**). Consistent with the fluorescence data, ManS-sfGFP increased the Man content of -insoluble (AKI) polymers after 9 h of induction, with no further increases at later timepoints (**Figure 1B**). In contrast, sfGFP-ManS required 24 h of induction to reach its maximal expression and still accumulated ≥2-fold less Man than ManS-sfGFP. Therefore, the C-terminal ManS tagged construct expressed well and provided a reliable proxy for the relative quantity of mannan made.

**Figure 1.**
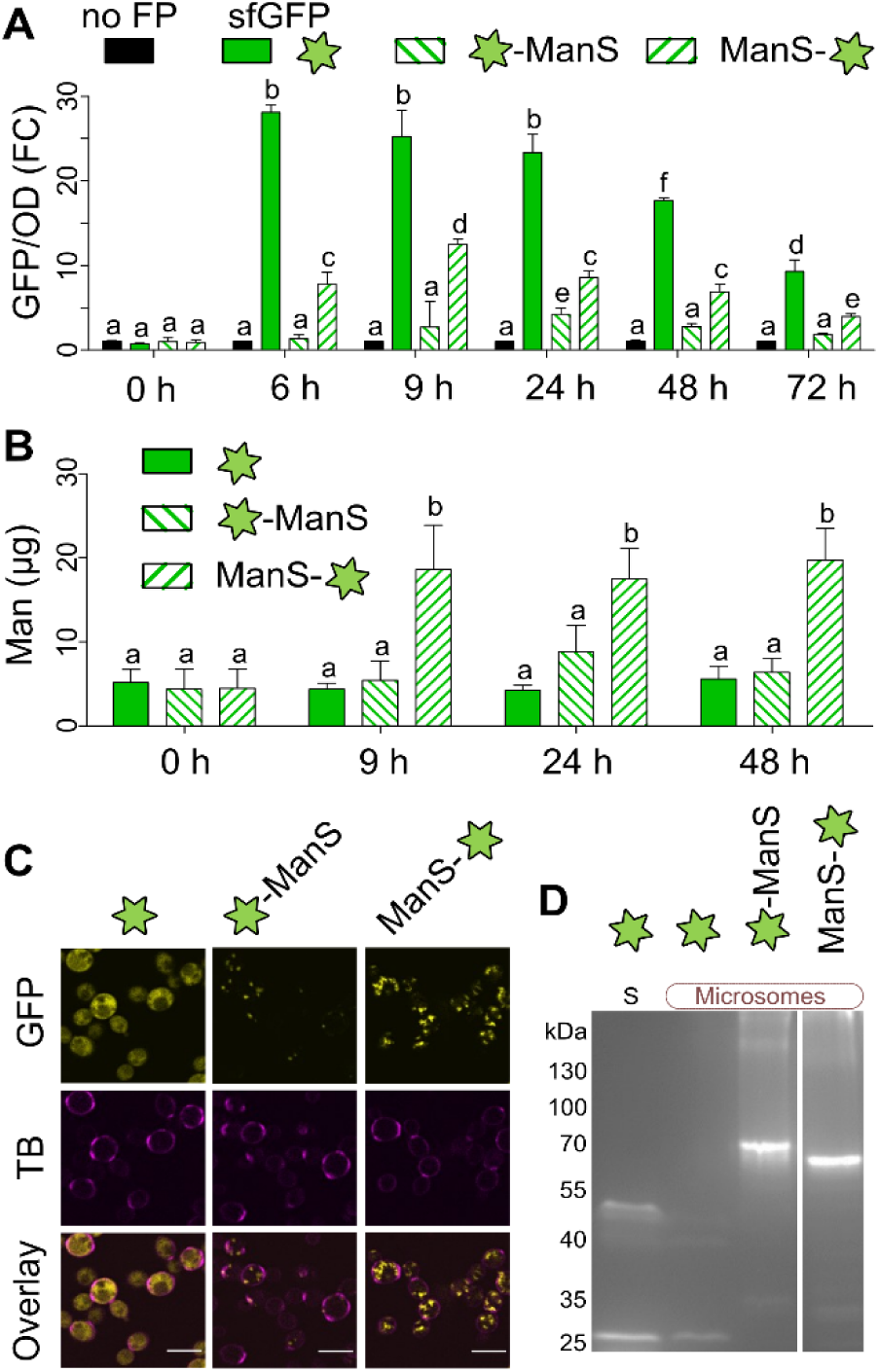
Expression of sfGFP tags in *Pichia*. Pre-cultured cells were induced using methanol. (A) Relative FP expression levels (GFP/OD600) were normalized to the ‘no FP’ control (set to have FC = 1) at each time point. (B) Man content of AKI polymers in equal aliquots at different timepoints. In (A) and (B), data show mean + SD from three biological replicates. Letters denote significant changes (one-way ANOVA with Tukey test, *P* < 0.05). (C) Protein localization at 24 h post-induction. Cell walls were counterstained with trypan blue (TB). Scale bars = 5 µm. (D) In-gel fluorescence of soluble (S) and microsomal proteins after electrophoresis in non-denaturing conditions.

Compared to the cytosolic distribution of free sfGFP (**Figure 1C**), tagged ManS localized in intracellular punctae, with brighter and a greater number of fluorescent bodies observed for the C-terminal sfGFP fusion. To ensure the sfGFP remains attached to ManS, soluble and microsomal protein fractions were examined for in-gel fluorescence (**Figure 1D**). On its own, sfGFP primarily showed the expected 27 kDa in the soluble fraction, with an additional ∼50 kDa band representing dimers. Tagged ManS proteins were only detected in the microsomal fraction, just below the predicted ∼87 kDa, with negligible signs of cytosolic fluorescence, sfGFP cleavage or significant protein glycosylation (**Figure 1D**). Therefore, the sfGFP-tagged ManS enzymes could be an enabling tool for future studies to decipher the detergent solubility, topology, and substrate specificities of CSL proteins.

In addition to sfGFP, which fluoresces regardless of its partner protein’s folding status,^13^ we found that Venus (a yellow FP that tolerates lower pH values) and mRuby2 (a monomeric red FP) express well in *Pichia* (**Figure S1A**). However, the three FPs had different dynamic ranges for fluorescence intensity in yeast. In the pPICZ B vector, ManS-Venus had ∼20x higher fluorescence compared to no FP control, followed by ManS-sfGFP (∼5x), while ManS-mRuby2 fluorescence was barely detected (∼1.25x) with a plate reader. Despite this large variation, all ManS-FP constructs produced more Man-containing polymers than sfGFP-ManS (**Figure S1B**) and the FP-only controls. Since the ManS-Venus construct had the highest dynamic range, we investigated if Glc or Man supplementation of YPM (yeast extract, peptone, methanol) medium could boost enzyme expression (**Figure S1C)** and cell wall synthesis (**Figure S1D**). Glycerol (the typical carbon source for *Pichia* ^12^) still performed best for mannan production, even though Glc and Man increased biomass and ManS expression compared to YPM alone. Overall, ManS-sfGFP produced the most glucomannan of the tested combinations, so we prioritized this construct and glycerol supplementation (YPM+G) for further *Pichia* experiments in this study.

### Golgi Localization and Modification of the Metabolic Sink for GDP-Man

We hypothesized that ManS-sfGFP elongated β-mannan in the Golgi apparatus, which was previously visualized in *Pichia* using ScGOS1 as a reporter protein.^14^ We assembled a ScGOS1-mRuby2 construct driven by the constitutive *pGAP*, and monitored co-expression with ManS-sfGFP after methanol induction (**Figure 2A**). ManS and the Golgi marker ScGOS1 localized in overlapping punctae (**Figure 2B**), with a largely consistent distribution at all tested timepoints. However, ManS-sfGFP expression, localization and function could be severely impaired by deleting its N-terminal region containing the first transmembrane domain (ΔTM1; **Figure S2A**). The ΔTM1-ManS-sfGFP expressed poorly (**Figure S2B)**, no longer produced mannan (**Figure S2C**), and aggregated in one or a few larger subcellular compartments per cell (**Figure S2D)**. The critical importance of the TM1 region for Golgi localization and protein folding or stability could explain the lower activity of the N-terminal tagged ManS construct (**Figure 1**) and the malfunction of prior CSLA domain swaps in this region.^12^

**Figure 2.**
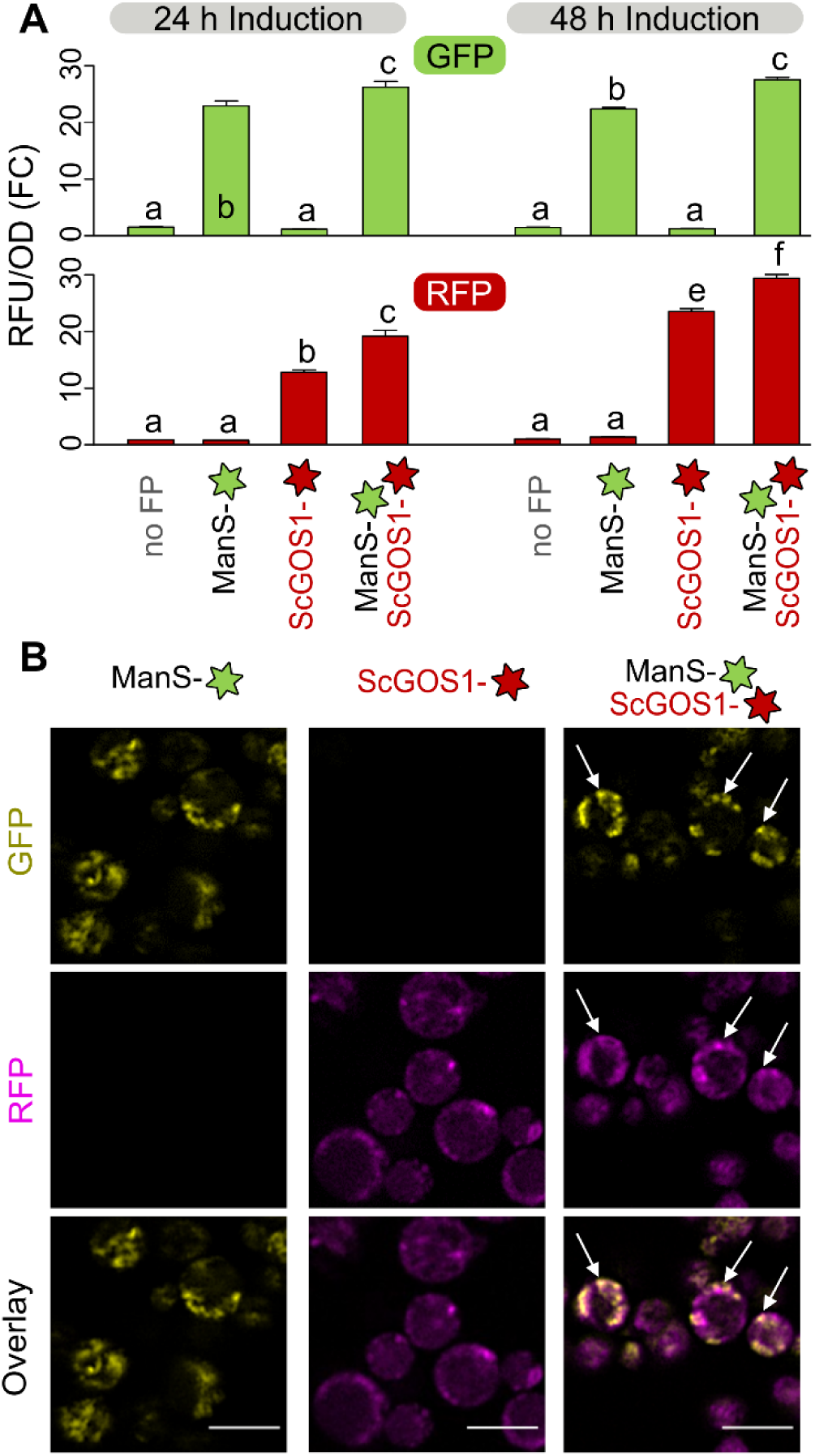
Co-localization of ManS with a Golgi marker. (A) Fold change (FC) in GFP/OD after methanol induction. The ‘no FP’ control was set to 1. Data show the mean + SD of three biological replicates. Distinct letters mark significant changes (one-way ANOVA with Tukey test, *P* < 0.05). (B) Arrows mark overlapping signals at 48 h. Scale bars = 5 µm.

To test whether plant β-mannan production is limited by competition for GDP-Man with yeast protein mannosylation, we introduced ManS-sfGFP in the SuperMan5 commercial strain that disrupts the *OCH1* α-1,6-mannosyltransferase.^15^ ManS-sfGFP generally had lower fluorescence in SuperMan5 compared to the X-33 wild-type (WT) background (**Figure 3A, Figure S3A**), but its localization was not altered (**Figure 3D)**. Since SuperMan5 restricts protein mannosylation, we hypothesized that this mutant strain would free up GDP-Man for plant glucomannan elongation. However, WT + ManS outperformed Superman5 + ManS in producing alcohol-insoluble Man-rich polymers (**Figure 3B**). After alkaline pre-treatment, the resulting AKI polymers were digested with an endo-β-mannanase to solubilize plant hemicellulose. ManS-sfGFP produced glucomannan with a similar composition (Glc:Man ratio of 1:3) in both X-33 and SuperMan5 backgrounds, despite a lower yield in the latter strain (**Figure 3C**). Similar trends in ManS-sfGFP expression (**Figure S3A**) and AKI composition (**Figure S3B**) were observed for two additional SuperMan5 strains that are deficient in protease activity. Although SuperMan5 was previously reported to have normal doubling time,^15^ our cultivation of this glycoengineered strain led to partial propidium iodide (PI) uptake even without plant ManS expression (**Figure 3D**), which can be toxic to *Pichia* cells.^12^ These results indicate that factors other than GDP-sugar allocation could be rate-limiting for ManS, since *OCH1* disruption did not promote the redistribution of Man units from yeast mannoproteins into plant glucomannan. Indeed, ManS activity in the X-33 background was previously elevated by co-expression with plant Mannan Synthesis-Related (MSR) proteins that act as co-factors.^8^

**Figure 3.**
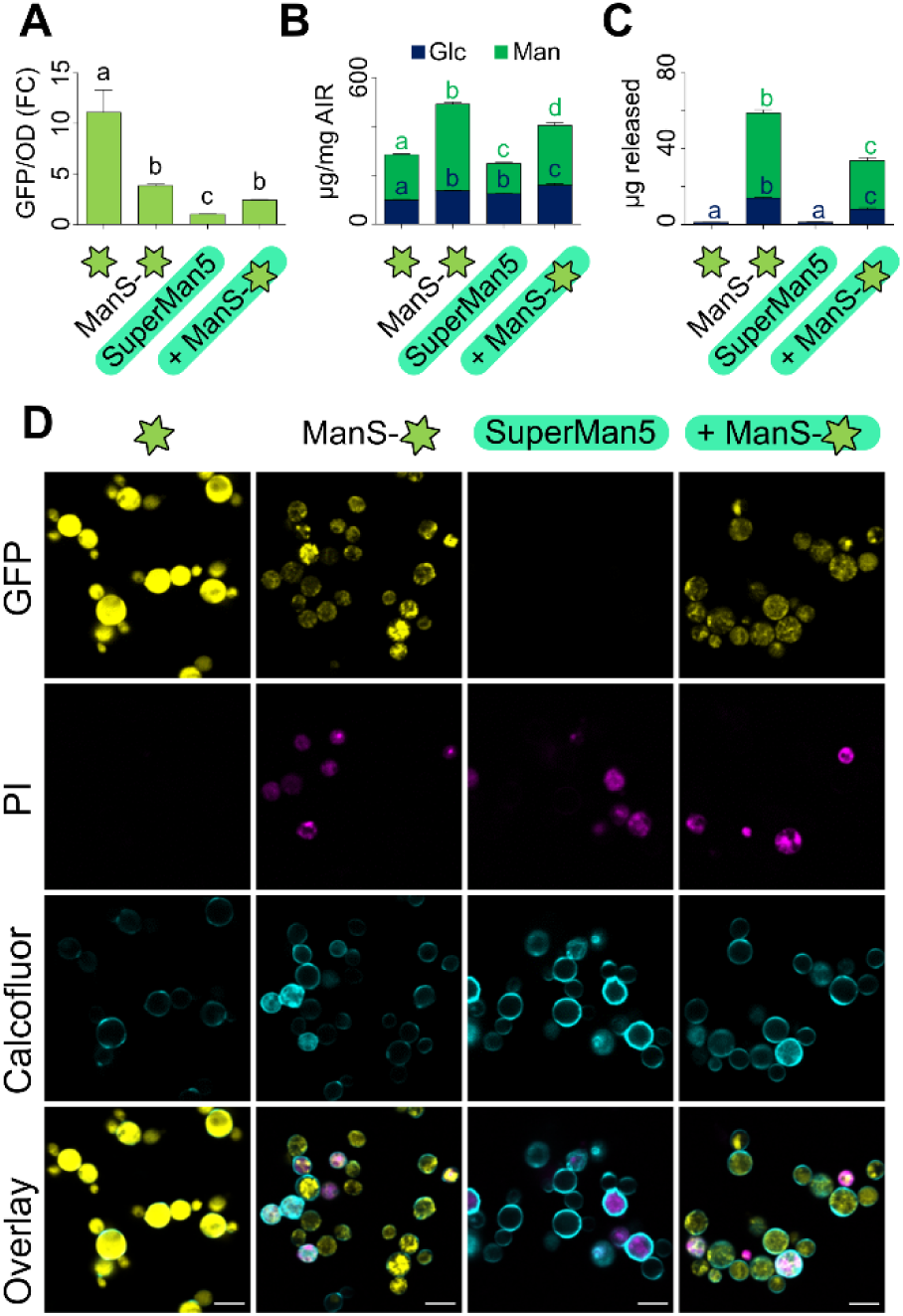
Glucomannan synthesis in the SuperMan5 strain. (A) Fold change in GFP/OD relative to SuperMan5 control after 48 h in YPM+G. (B) Composition of total alcohol-insoluble residue (AIR) and (C) AKI carbohydrates solubilized by endo-β-mannanase digestion. Data show the mean + SD of six (A and B) or four (C) biological replicates. Different letters denote significant changes (one-way ANOVA with Tukey test, *P* < 0.05). (D) Confocal images of cell permeable to PI. Calcofluor was used as a counterstain. Scale bars = 5 µm.

### Chemical Treatments of ManS-Expressing Cells

Using the fluorescent strains established in this study and known disruptors of cell wall glycans or their secretion, we evaluated how exogenous treatments modulate yeast growth, morphology, and polysaccharide biosynthesis. ManS-sfGFP expression and products were quantified in methanol-containing media supplemented with dimethyl sulfoxide (DMSO) only, Zymolyase 20T, Anidulafungin, or Brefeldin A (BFA). With either DMSO or BFA, ManS-expressing strains accumulated more Man but less Glc than the empty vector and ScGOS1 Golgi marker alone, consistent with a growth penalty for glucomannan production (**Figure 4A**). BFA, which induces intracellular aggregates in plants,^16^ led to more diffuse ManS-sfGFP localization (**Figure 4C**), but still resembled the glycan profile of the DMSO control. In contrast, Zymolyase and Anidulafungin dramatically reduced yeast biomass and the insoluble polysaccharide content for all genotypes. Zymolyase 20T directly hydrolyzes fungal β-1,3-glucans in the extracellular matrix, while Anidulafungin disrupts the same polymers via a distinct mode of action. As an antifungal drug that directly binds and inhibits glucan synthases with multiple TM domains,^17^ Anidulafungin led to severe swelling (**Figure 4C**), increasing cell size by up to 3-fold compared to the DMSO control. Although ManS-sfGFP sig nificantly increased the % Man of AKI in all four treatments (**Figure 4B**), Anidulafungin-treated cells were the least productive in terms of total carbohydrate content (**Figure 4A**). The loss of sfGFP fluorescence in swollen Anidulafungin-treated cells could be due to severely impaired Golgi membrane integrity and/or ManS stability (**Figure 4C**). In contrast, *Pichia* cells with partly inactivated synthases were recently shown to have milder 17–28% reductions in β-1,3-glucans, but increased mannoprotein content and GFP expression.^18^ Similar to our findings, the genetic impairment of glucans increased yeast cell size.^18^ In contrast, the Zymolyase treatment did not alter the punctate ManS-sfGFP localization even in cells with lysed walls and reduced labeling with calcofluor white (arrows in **Figure 4C**). The pharmacological results underscore the potential to redesign *Pichia* cell walls and to potentially displace fungal components with plant-based polymers. Our data indicate that extracellular digestion could be a superior strategy to preserve membrane integrity (**Figure 4B**).

**Figure 4.**
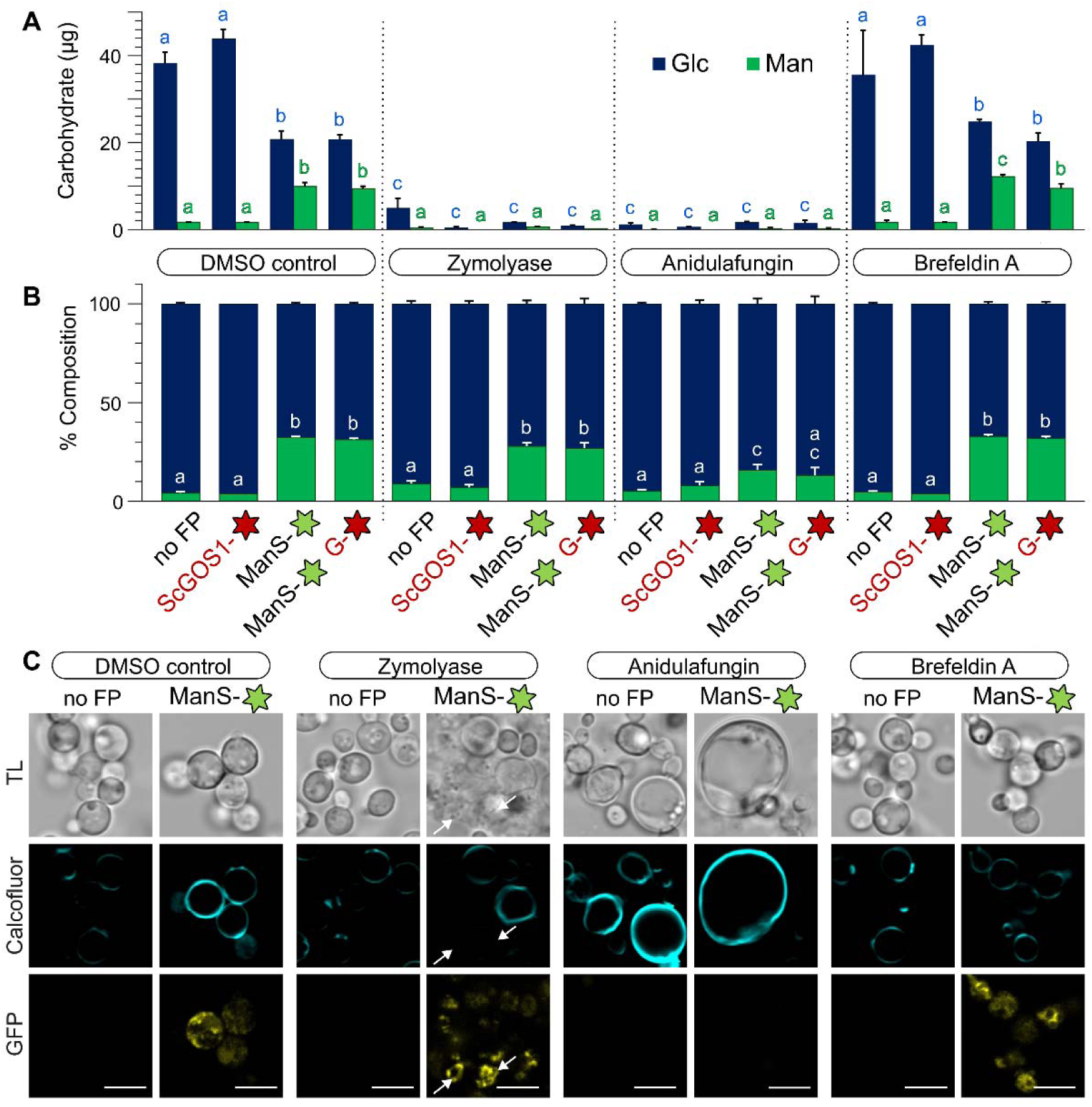
Modulating yeast wall composition and morphology using exogenous treatments. Cells pre-cultured in YPD were transferred for 48 h with YPM plus the indicated treatments. Absolute amounts (A) and relative composition (%, w/w) of monosaccharides in AKI polymers (B). Bars show mean + SD of three biological replicates per genotype. G is an abbreviation for ScGOS1. Letters denote significant differences obtained by one-way ANOVA with Tukey’s pairwise (*P* < 0.05). (C) Confocal images of treated *Pichia* cells counterstained with calcofluor white. Similar GFP signals were observed with or without the expression of ScGOS1. Arrows mark cells with Zymolyase-digested walls but unaffected ManS-sfGFP localization. Scale bars = 5 µm.

### Yeast Systems to Accelerate the Prototyping of Plant Cell Wall Synthases

Finally, we applied ManS and sfGFP to test alternative expression vectors that could speed up the prototyping of cell wall-related enzymes for downstream applications in plant synthetic biology. While GoldenPiCS vectors were previously used to assemble CSLA and CSLC chimeric enzymes^12^ and offer well-characterized *Pichia* regulatory elements^19^ for multi-gene constructs, they are not compatible with the Golden Gate fusion site syntax established in the plant research community.^20^ We therefore tested new level 1 (L1) episomal vectors^21^ (*pPAP002* for *Pichia* and *pAGT572_Nemo* for *Saccharomyces cerevisiae*) that are compatible with Plant Modular Cloning (MoClo) standards. While L1 episomal plasmids require antibiotic or auxotrophic selection during yeast protein induction, they offer high transformation efficiencies using circular DNA compared to vectors for genome integration. To our initial surprise, *pPAP002 + ManS* grew poorly and had 97% lower AKI biomass than *pPAP002* + *sfGFP* in YPM + G medium (**Figure 5A**). Since the methanol-inducible *pCAT1* promoter in *pPAP002* could be de-repressed by glucose or glycerol depletion,^22^ we generated a new *Pichia* L1 episomal vector (*L1_2F_ePH*) that can accept any promoter. Whether *Pichia* cells were fed methanol, limited glucose or both, *pCAT1*:*sfGFP* showed a growth penalty (**Figure S4A**) and at least 4x higher fluorescence than *pAOX1:sfGFP* (**Figure S4B)**. The *pAOX1* promoter was better for ManS activity in *Pichia* episomal expression (**Figure 5B**), even though the original integrative vector provided the highest Man content. Similarly, we discovered that *S. cerevisiae* cells produced Man-rich polymers via episomal ManS-sfGFP expression (**Figure 5C**), only if the yeast biomass accumulated before galactose induction of the strong *pGAL1* promoter (**Figure S4C–F**). Although *Pichia* produced more plant mannan, *S. cerevisiae* had a lower background level of endogenous Man in the AKI fraction and thus offers more room for future efforts to improve plant CSL enzyme activity via artificial intelligence (AI) protein algorithms and/or continuous directed evolution.^23^

**Figure 5.**
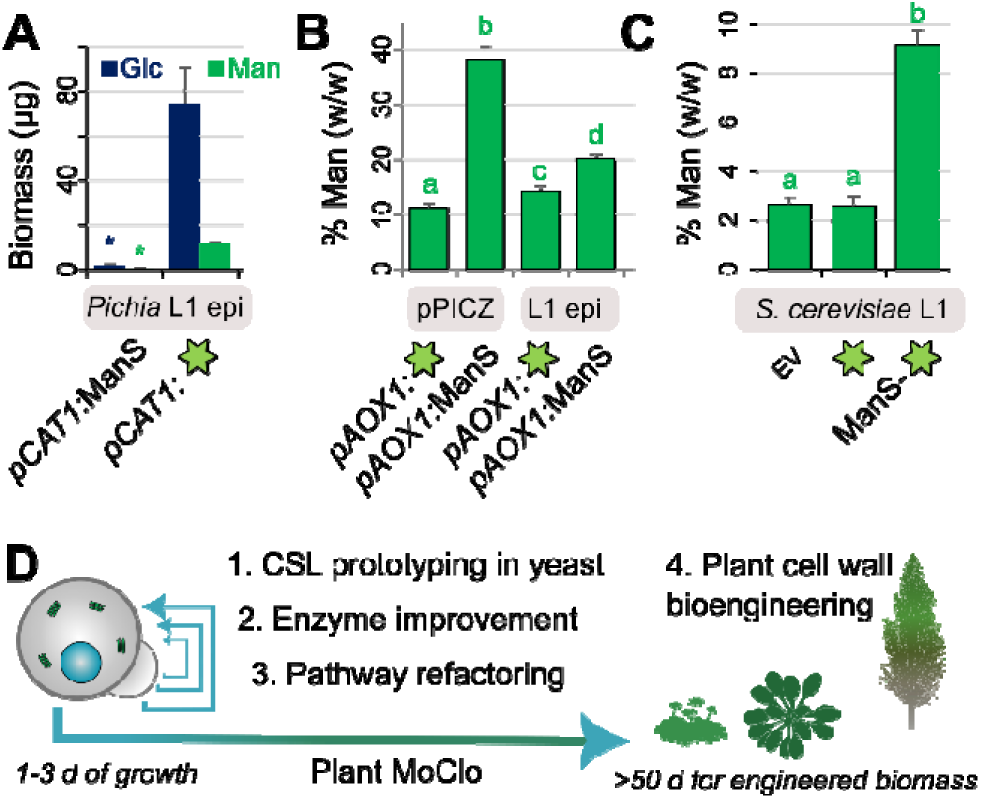
Cell wall prototyping using yeast episomal vectors compatible with Plant MoClo. (A) Absolute AKI composition of *Pichia* cells with a *pCAT1*-driven L1 episomal vector, after growth in YPM + G. (B) Relative Man level in *Pichia* AKI after growth on YPM + G using integrative and episomal vectors driven by *pAOX1*. (C) Relative Man level in *S. cerevisiae* AKI using an L1 episomal vector, after biomass accumulation and galactose induction. (D) Strategy to screen and improve CSL enzymes and related glycan pathways in yeast using Plant MoClo-compatible L1 vectors. Yeast data would help to prioritize constructs for stable plant transformation (e.g. *Marchantia*, *Arabidopsis*, or *Populus* species).

## CONCLUSIONS

In this study, we bioengineered yeast cells via chromosomal or episomal constructs to synthesize hemicellulose within 9 to 24 h of ManS induction. Our FP-tagged design illuminated the ManS subcellular localization and relative expression to probe new avenues for cell wall improvement. Using Plant MoClo-compatible vectors, a ManS-FP mutant library could be screened for improved expression and polysaccharide synthesis after 3 days of yeast growth, before waiting 2-3 months to generate stable plant transformants for the most promising variants (**Figure 5D**). By showing that *S. cerevisiae* is a suitable host for plant mannan synthesis,^7^ future biomaterials research will benefit from the availability of a genome-wide collection of yeast mutants and advanced tools (e.g. OrthoRep) to improve plant enzymes in yeast for ultimate deployment in crops through genome editing.^23^ To address the metabolic bottlenecks such as increasing hexose sugar content in lignocellulosic biomass, future studies could test the roles of sugar transporters and re-examine the CSL enzyme topology around the Golgi membrane.^24^ Although *in vitro* yeast experiments predicted that dedicated transporters are required to deliver GDP-sugars for ManS activity in the Golgi lumen,^25^ none of the canonical Arabidopsis GDP-Man transporters affected glucomannan synthesis *in planta.*^26^ Yeast prototypes offer *in vivo* platforms to “fail fast” when engineering steps involved in cell wall synthesis^27^ or fiber degradation,^28^ and thus de-risk strategies to test in plants. Since the L1 plasmids are limited to single transcriptional units, additional *Pichia* and *Saccharomyces* vectors compatible with Plant MoClo^29^ are still needed to evaluate multi-gene combinations.

## METHODS

### Vector construction and transformation

**Table S1** summarizes the plasmids used in this study and their molecular cloning strategies. The pPICZ classical cloning and GoldenPiCS assembly into the BB3aZ_14 vector^19^ were previously described.^8,12^ Unless otherwise indicated, all *Pichia* ManS constructs were driven by the methanol-inducible *pAOX1*, integrated into the *AOX1* genomic region, and selected using Zeocin. New parts were amplified with the High-Fidelity Phusion DNA Polymerase (Thermo Fisher Scientific, Cat# F530L) and the primers listed in **Table S2**. The *BB3rN_14* backbone^19^ was used for the *pGAP:ScGOS1* expression and Nourseothricin selection. The *pPAP002* vector^21^ was modified using NEBuilder HiFi DNA Assembly Cloning Kit (New England Biolabs) to yield a *Pichia* episomal expression *L1_2F_ePH* vector that can accept any Promoter-Gene-Terminator combination following the Plant MoClo syntax.^30^ *Saccharomyces* episomal expression vectors were assembled using *pAGT572_Nemo* backbone.^21^ All plasmids were cloned *in silico* using the Assembly function in Benchling (https://benchling.com) to inspect the design outcomes (e.g. in-frame fusions with the fluorescent proteins). Sanger sequencing was used to verify all new parts that were cloned using PCR amplicons. A key advantage of Golden Gate cloning (e.g. with GoldenPiCS or Plant MoClo vectors) compared to other methods is the unparalleled fidelity to yield the desired product.^29^ Whole plasmid sequencing performed by Plasmidsaurus using Oxford Nanopore Technology showed that positively genotyped vectors contained the desired sequences. Linearized integrative plasmids (*pPICZ B* and GoldenPiCS vectors) and circular episomal plasmids were transformed into *Pichia pastoris* (X-33 wild-type, unless otherwise stated) via electroporation.^31^ The *pAGT572_Nemo* episomal constructs were transformed into *Saccharomyces cerevisiae* BY4742 using Frozen-EZ Yeast Transformation II Kit (Zymo Research, # T2001). The GoldenPiCS Kit (#1000000133; donated by the Gasser/Mattanovich/Sauer group), and Plant MoClo-compatible vectors (MoClo Toolkit, #1000000044; *pPAP002*, #153489; and *pAGT572*_*Nemo*, #153487; all donated by Marillonnet Lab) were purchased from AddGene. SuperMan5 strains, belonging to the GlycoSwitch collection, were purchased with an academic license from Research Corporation Technologies (RCT; https://pichia.com/). After growth on agar plates containing appropriate selection media, at least three PCR-verified independent colonies with the desired constructs were screened in liquid cultures.

### Yeast growth and medium composition

Cells were cultivated in stackable shaking incubators (Thermo Fisher MaxQ 6000 or Eppendorf Innova 42R) at 30°C and 250 rpm in 24-well plates, for 48 h unless otherwise specified. Most *Pichia* batches were grown in YP base medium containing 1% (w/v) yeast extract, 2% (w/v) peptone was supplemented with 2.0% (w/v) dextrose/glucose for YPD; or 1.5% (v/v) methanol for YPM, or in YP containing more than one carbon source as specified in the figure legends. For **Figure S1** (A and B), buffered media were prepared according to the EasySelect *Pichia* Expression Kit (ThermoFisher Scientific). For *Pichia* episomal vectors, the liquid media was supplemented with hygromycin (150 µg/mL) to ensure the plasmids were retained.

For *Pichia* biomass accumulation, cells were pre-cultured in ≥2 mL of glycerol-rich medium for at least 24 h followed by centrifugation at 2000 g for 2 min, and resuspension in a similar volume of methanol-containing medium for 24–72 h of induction. For the antifungal treatments, 2 mg of Zymolyase 20T (Carl Roth, Cat# 9324.3), 6 µg of Anidulafungin (Sigma Aldrich, Cat# SML2288-5MG), or 8 µg of Brefeldin A (VWR International GmbH, Cat# CAYM11861-10) dissolved in DMSO were applied per mL of *Pichia* culture. To isolate sufficient protein for in-gel fluorescence, we cultivated yeast for 24 h in 50 mL of YPD from an initial OD600 of 0.3, followed by centrifugation (5 min at 2000 *g*), resuspension, and growth in 50 mL of YPM for another 24 h.

For *Saccharomyces* biomass accumulation, cells were grown in 3 mL in YPD for 48 h followed by centrifugation at 2000 g for 2 min. Cells were resuspended in 3 mL of DO^Ura-^ (yeast drop out medium, excluding uracil to retain the *pAGT572_Nemo* vector) supplemented with 2% (w/v) galactose to induce protein expression for 24 h.

### Plate reader measurements and in-gel fluorescence

For plate reader assays, equal aliquots of cell suspensions were diluted with water (generally 1:10) to avoid saturation and analyzed in 96-well plates. OD600 and fluorescence (excitation/emission peaks: 485/511 for GFP, 513/527 nm for Venus, and 569/593 nm for mRuby2) were measured using Tecan M1000 (Grödig, AT) or BioTek Synergy H1 plate reader (Agilent Technologies, USA) plate readers. We compared relative fluorescent units (normalized to OD600) or fold changes relative to cells without FPs.

Microsomal proteins were extracted from *Pichia* cultures based on a prior method,^32^ with slight modifications, starting with a cell lysis containing 50 mM Tris-HCl (pH 7.6), 150 mM NaCl, and 1x Pierce protease inhibitors (Thermo Fisher, Cat# A32955). *Pichia* cells were milled with glass beads in a Retsch MM400 homogenizer for a total of 7 cycles, each consisting of 3 min at 20 Hz for 3 min, followed by cooling on ice. After centrifugation at 3000 *g* and 4°C for 20 min, the supernatant was transferred to new tubes and centrifuged at 20000 *g* for 1 h at 4°C to separate the microsomal membrane and soluble cytosolic fractions. Following a Bradford assay,^33^ 20 μg of proteins were diluted 1:5 with 5x Laemmli protein buffer and incubated for 5 min at 55°C. Samples were loaded alongside 2 μL of PageRuler ladder (Thermo Scientific) into a Mini PROTEAN Tetra cell (Bio-Rad, Hercules, US) with 1.0 mm thick gels, for electrophoresis in 1x ROTIPHORESE SDS-PAGE buffer (Carl Roth) at 100–150 V for 2 h. Protein were visualized (blue LED light plus GFP filter) in a UVP GelStudio PLUS Touch (Analytik Jena GmbH, DE).

### Confocal microscopy

Cells were stained with 0.01% (w/v) of the specified dye as previously described.^12^ Zeiss LSM880 micrographs (**Figures 1 and3**) were acquired using a 40x/1.2 water-immersion objective, a beam splitter MBS 488/561 and AiryScan mode. The excitation/emission peaks were 488/523 nm for GFP/YFP, 561/579 nm for RFP/trypan blue. In the remaining micrographs, Zeiss LSM900 images were acquired using a 63x/1.20 water-immersion objective with AiryScan mode and the following excitation/emission parameters: calcofluor (405/410–490 nm, SP545), GFP/YFP (488 nm/490–560 nm), propidium iodide (639 nm, 656–700 nm, LP655). For each panel, images were uniformly processed with ImageJ.^34^

### Yeast cell wall analyses

*Pichia* and *Saccharomyces* cells were lysed, washed, and dried to obtain cell wall AIR or to directly enrich glucomannan AKI polymers as previously described.^8^ For monosaccharide analyses, 50 μL aliquots of the AKI polymers suspension or 300 μg of dried AIR material were hydrolyzed with 2 M trifluoroacetic acid, before drying and re-suspension in 400 µL of 30 µg/mL Ribose. For Figure 3C, glucomannan in 50 μL AKI aliquots was solubilized as previously described^12^ but with 0.1 U of endo-1,4-β-mannanase (Megazyme, E-BMANN), prior to acid hydrolysis and monosaccharide composition analysis. Samples and monosaccharides standards were separated and quantified using high-performance anion-exchange chromatography coupled with pulsed electrochemical detection (HPAEC-PAD). After injecting 10 µL of each sample/standard, the HPAEC-PAD setup applied a previously described 30 min eluent gradient^12^ on a Metrohm 940 Professional IC Vario system equipped with a Metrosep Carb 2–250/4.0 column.

## ASSOCIATED CONTENT

**Supporting Information**. A single PDF document containing the following figures and tables is available free of charge.

**Figure S1.** *Pichia* expressing ManS with or without C-terminal FPs.

**Figure S2.** Impact of N-terminal region on ManS-sfGFP expression and activity.

**Figure S3.** ManS-sfGFP performance in three SuperMan5 strains.

**Figure S4.** Feeding strategies and episomal vector performance in two yeast species.

**Table S1**. Yeast strains and vectors used in this study.

**Table S2**. Primers used for cloning and genotyping.

**Supplemental References**

## ACKNOWLEDGMENT

C.V. thanks Benita Schmitz and Christine Wagner for technical assistance, and gratefully acknowledges UF/IFAS startup funding, and the USDA National Institute of Food and Agriculture, Research Capacity Fund (Hatch) project 7004470, the UF Horticultural Sciences department. This work was also in part supported by the Deutsche Forschungsgemeinschaft (DFG; German Research Foundation grant no. 414353267 to C.V.) and by core funding (Leibniz Association) from the Federal Republic of Germany and the state of Saxony-Anhalt.

## ABBREVIATIONS

ManS: β-mannan synthase
AKI: alkaline-insoluble polysaccharides

## Author Contributions

C.V. designed the experiments. A.G.-O., M.R., M.Q.. S.C. and C.V. collected and analyzed data.

C.V. prepared and revised the manuscript using draft text and figures from all authors.

