## Supporting Information for "Illuminating Glucomannan Synthases to Explore Cell Wall Synthesis Bottlenecks"

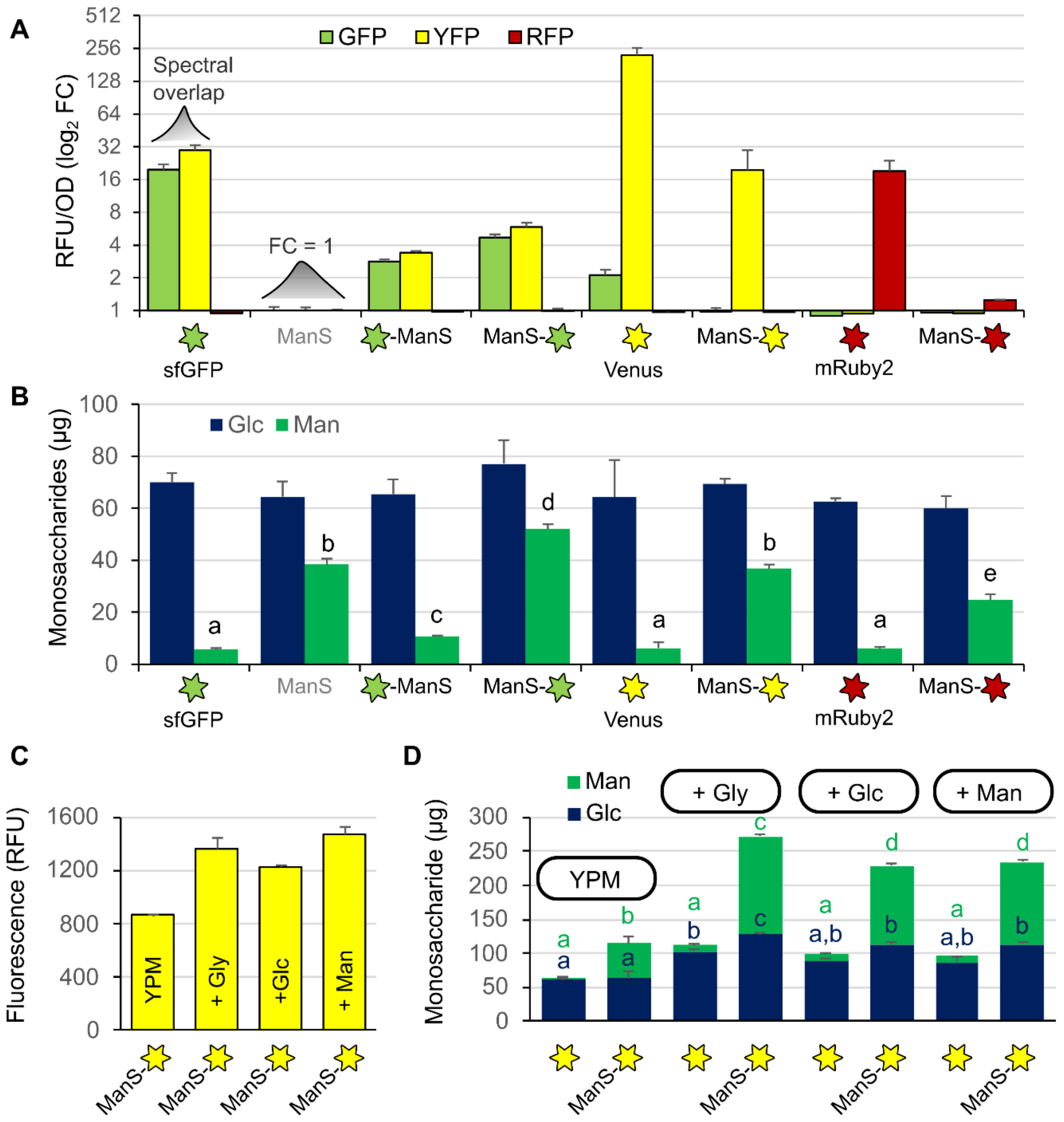


**Figure S1.** *Pichia* expressing ManS with or without C-terminal FPs. (A) Log2 fold changes (FC) in relative fluorescence units normalized to OD600 biomass. FC was set to 1 for the untagged ManS control in each of the three fluorescence channels that were analyzed with a plate reader. GFP and YFP channels show spectral overlap, particularly for the sfGFP tag. Yeast strains were cultivated for 24 h in BMGY medium for biomass production followed by 24 h growth in BMMY. (B) Composition of AKI wall polymers from samples in (A). (C) Relative fluorescence of ManS-Venus in YPM medium alone, or in YPM medium supplemented with 0.5% w/v Glycerol (Gly), Glucose (Glc) or Mannose (Man). (D) Composition of yeast AKI wall polymers from Venus and ManS-Venus lines after the feeding strategies shown in (C). For all panels, data show the mean + SD of at least 3 biological replicates and letters denote for significant changes (one-way ANOVA with Tukey test, $P < 0.05$).

**Supporting Information**

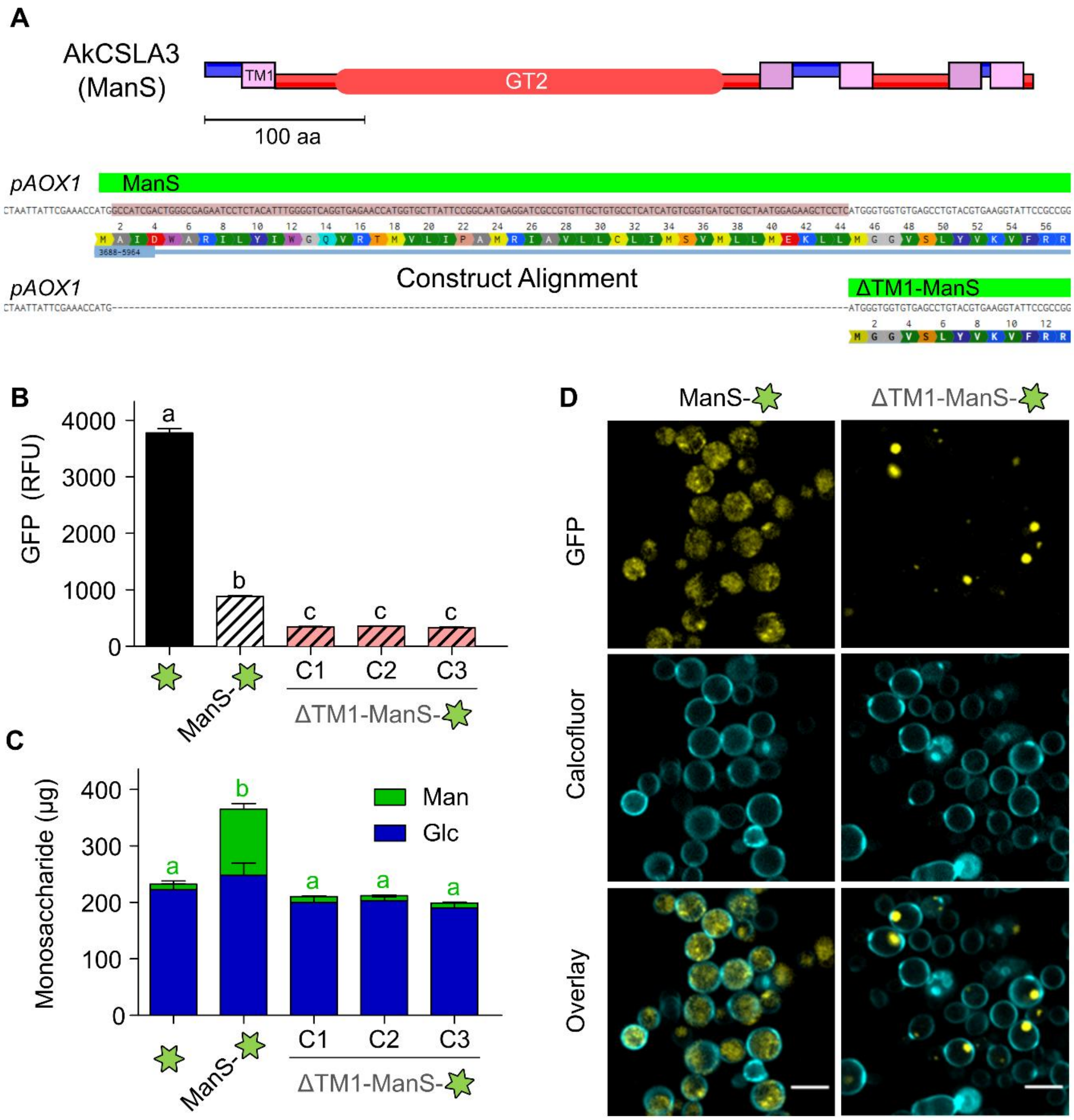


**Figure S2.** Impact of N-terminal region on ManS-sfGFP expression and activity. (A) Topology of AkCSLA3 (ManS in this study) adapted from an earlier publication.[1] Amino acids (aa) 24 to 44 are predicted to encode the first transmembrane domain (TM1). A plasmid sequence alignment from Benchling shows the *pAOX1* promoter followed by the ManS coding sequence. (B) Relative fluorescence units (GFP/OD) of sfGFP-tagged constructs using BB3_aZ integrative vectors and (C) composition of AKI wall polymers. *Pichia* cultures were cultivated in YPM + G for 48 h. Data show the mean + SD of 3 biological replicates and different letters denote significant changes (one-way ANOVA with Tukey test, $P < 0.05$). For the TM1 deletion constructs, three independent colonies (C1 to C3) were analyzed. (D) Localization of sfGFP-tagged ManS proteins with or without the N-terminal region. *Pichia* cell walls were counterstained with calcofluor. Scale bars: 5 µm.

### Supporting Information

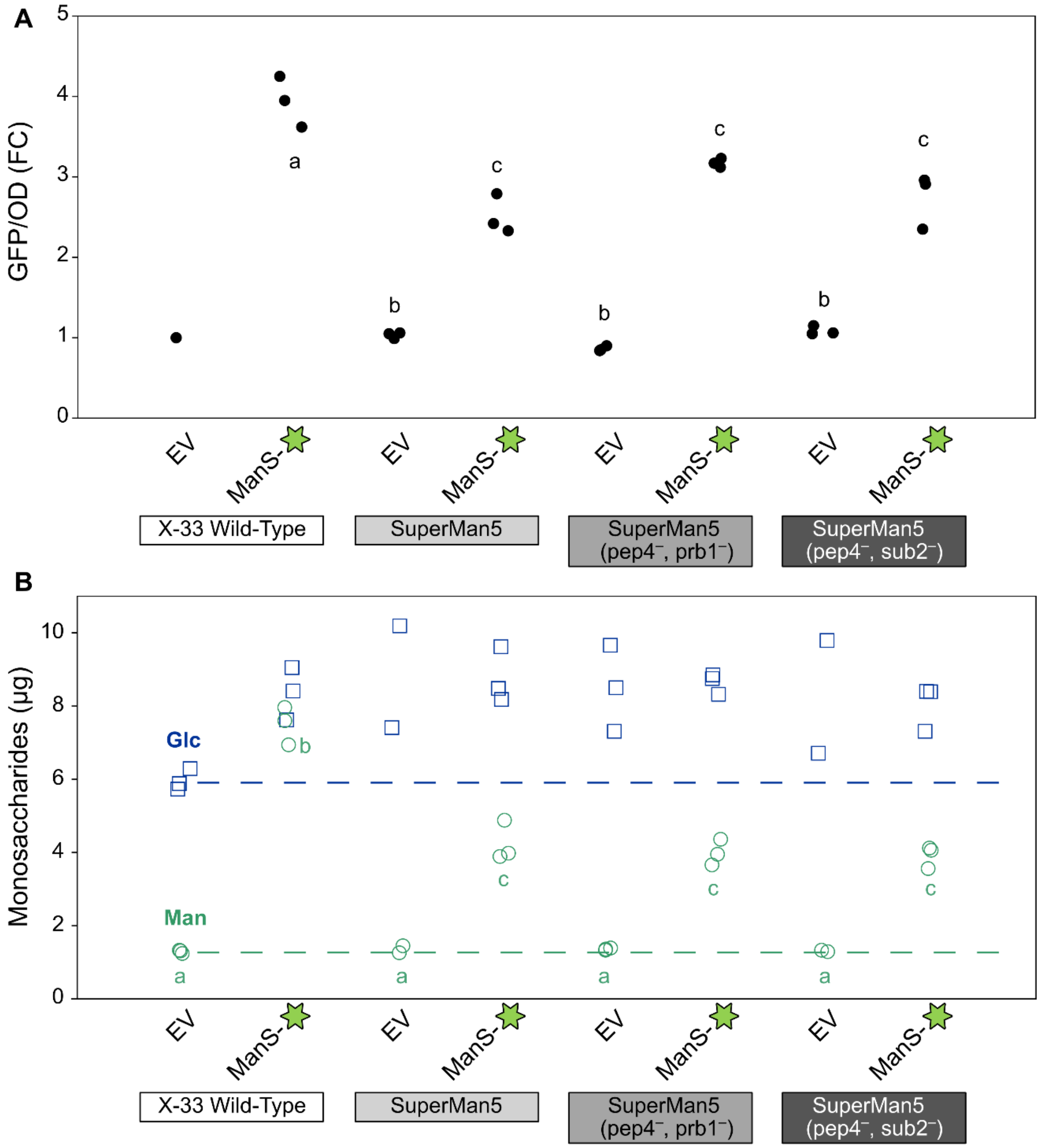


**Figure S3.** ManS-sfGFP performance in three SuperMan5 strains. (A) Fold changes (FC) in fluorescence (GFP/OD600) were normalized to pPICZ B empty vector (EV) control, which was set to have FC = 1 in the X-33 background after growth for 42 h in BMMY. The Superman5 (pep4⁻, prb1⁻) and Superman5 (pep4⁻, sub2⁻) strains are protease-deficient. (B) Monosaccharide composition of AKI polymers in equal aliquots. Graphs show independent colonies for each vector in four different *Pichia* backgrounds. Letters in (B) denote significant changes in Man levels (one-way ANOVA with Tukey test, $P < 0.01$).

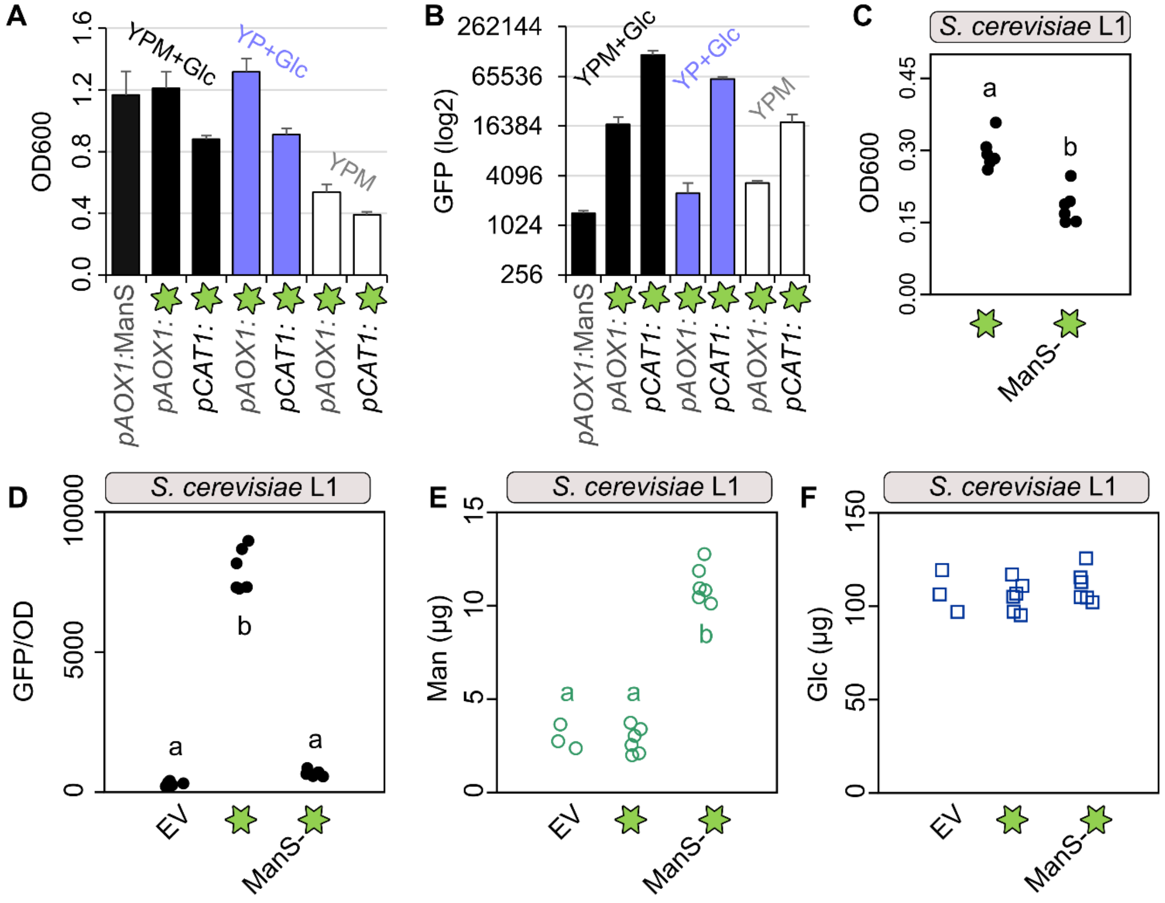


**Figure S4.** Feeding strategies and episomal vector performance in two yeast species. (A) Optical density and (B) GFP fluorescence (logarithmic scale) of *pAOX1* versus *pCAT1* episomal *Pichia* strains after 48 h of growth in YP-based media supplemented with 2% v/v methanol (M) and/or 0.5% w/v Glc. The *pAOX1:ManS* episomal construct is untagged and serves as a negative control for fluorescence. Bars show mean + SD of three biological replicates per genotype. (C) Optical density of *Saccharomyces cerevisiae* after 24 h of direct cultivation in DO$^{Ura-}$ supplemented with 2% (w/v) galactose to induce protein expression of an L1 episomal vector. (D) Relative fluorescence units of *S. cerevisiae* L1 empty vector (EV), sfGFP and ManS-sfGFP after biomass accumulation in YPD for 48 h, followed by 24 h of induction in DO$^{Ura-}$ with 2% (w/v) galactose. This two-step cultivation strategy (corresponding to Figure 5C) produces plant mannan (E) without compromising biomass accumulation (F), as shown by consistent levels of yeast glucans. In (C) to (F), dots represent independent colonies and letters mark significant differences obtained by one-way ANOVA with Tukey's pairwise ($P < 0.05$).

### Supporting Information

**Table S1**. Yeast strains and vectors used in this study. For C-terminal fusions in the pPICZ B vector (EasySelect Pichia Expression kit, ThermoFisher Scientific), the fluorescent proteins (FPs) were inserted using Eco72I and XhoI sites and classical restriction enzyme (RE) cloning. The ManS coding sequence was trimmed to remove its stop codon and ensure in-frame FP fusion using Bsp119I and Eco72I RE cloning in pPICZ B + FP vectors. Sequences were domesticated and assembled into GoldenPiCS vectors, as directed by the original publication[2], using Golden Gate cloning. *Pichia* episomal vectors were assembled using *pPAP002* or a new *L1_2F_ePH* vector modified from *pPAP002* to accept any MoClo promoter-terminator combination. For *Saccharomyces* cerevisiae expression, genes were assembled in the L1 episomal vector *pAGT572_Nemo*,[3] containing a galactose-inducible promoter *pGAL1*.

| **Yeast strains** and plasmids | **Reference** | **Assembly** | **Promoter** |
|---|---|---|---|
| ***Pichia pastoris X-33*** | EasySelect Pichia kit | - | - |
| ***Pichia pastoris* SuperMan5 strains** | RCT's Pichia Glycoswitch[4] | - | - |
| *pPICZ B* | EasySelect Pichia kit | RE cloning | Inducible, pAOX1 |
| *pPICZ B + sfGFP* | prior work[5] | | |
| *pPICZ B + sfGFP-ManS* | prior work[5] | | |
| *pPICZ B + ManS-sfGFP* | this study | | |
| *pPICZ B + Venus* | this study | | |
| *pPICZ B + ManS-Venus* | this study | | |
| *pPICZ X + mRuby2* | this study | | |
| *pPICZ X + ManS-mRuby2* | this study | | |
| *BB3aZ_14 + pAOX1:ManS-sfGFP:RPP1Btt* | prior work[1] | GoldenPiCS | Inducible |
| *BB3aZ_14 + pAOX1:ΔTM1-ManS:sfGFP-RPP1Btt* | this study | | Inducible |
| *BB3rN_14 + pGAP:ScGOS1-mRuby2:ScCYC1tt* | this study | | Constitutive |
| *pPAP002* | prior work[3] | Plant MoClo | Inducible, pCAT1 |
| *pPAP002 + sfGFP* | this study | | |
| *pPAP002 + ManS* | this study | | |
| *L1_2F_ePH* | this study | | any promoter |
| *L1_2F_ePH + pAOX1:sfGFP:RPP1Btt* | this study | | Inducible, pAOX1 |
| *L1_2F_ePH + pAOX1:ManS:RPP1Btt* | this study | | |
| ***Saccharomyces cerevisiae BY4742*** | prior work[6] | - | - |
| *pAGT572_Nemo* | prior work[3] | Plant MoClo | Inducible pGAL1 |
| *pAGT572_Nemo + sfGFP* | this study | | |
| *pAGT572_Nemo + ManS-sfGFP* | this study | | |

### Supporting Information

**Table S2**. Primers used for cloning and genotyping. Restriction enzyme recognition sites are in bold and Golden Gate fusion sites are underlined. The insertion sites in GoldenPiCS vectors are flanked by commonly used M13 sequencing/genotyping primers. The pPICZ B + ManS-sfGFP was cloned by amplifying an unpublished pPICZ B + AkCSLA3-2A-sfGFP plasmid using 5' Phosphorylated, HPLC-purified primers, and circularizing the resulting amplicon.

| Target and Application | Forward Primer | Reverse Primer |
|---|---|---|
| pPICZ B + ManS-sfGFP | [5' Phos] TTTTCACTAGGAACAAAGGTGCCA | [5' Phos] atgcgtaaaggcgaagagctg |
| C-ter Venus for pPICZ | caa**CACGTG**tcatgtctaaaggtgaagaattattcactgg | gac**CTCGAGTC**Atttgtacaattcatccataccatg |
| ManS for C-ter fusions in pPICZ | a**TTCGAA**acgatatggccatcgactg | caa**CACGTG**tcatgtctaaaggtgaagaattattcactgg |
| C-ter mRuby2 for pPICZ | caa**CACGTG**tcatggtgtccaaaggagagga | gtg**CTCGAG**ctacttatacaattcatccataccaccg |
| Venus for BB1 | cgct**GGTCTC**aCATGtctaaaggtgaagaattattc | acga**GGTCTC**gaagcttatttgtacaattcatccatacc |
| mRuby2 for BB1 | atct**GGTCTC**aCATGgtgtccaaaggagaggag | acga**GGTCTC**gaagcttacttatacaattcatccataccaccg |
| Fs2-ScGOS1-new for BB1_23 | acga**GGTCTC**aCATGAGCTCACAACCGTCTTTCG | gctc**GGTCTC**aCTCCCCATGTGAAAAACAAAAACAG |
| new-mRuby2-Fs3 for BB1_23 | atat**GGTCTC**aGGAGTGTCCAAAGGAGAGGAGTTA | acga**GGTCTC**gaagcTTActtatacaattcatccataccaccg |
| Fs3-sfGFP-new | atat**GGTCTC**aGCTTcccgtaaaggcgaagagctg | taat**GGTCTC**actatttgtacagttcatccataccatgc |
| new-RPP1Btt-Fs4 | atat**GGTCTC**aatagggatgatactttaattgatgc | tcga**GGTCTC**cAGCGgctcctctagataccatcaag |
| pAOX1 for L0_12 | ttc**GAAGAC**tcGGAGagacgaaaggttgaatgaaac | cgc**GAAGAC**gccattgtttcgaataattagttgtttttg |
| RPP1Btt for L0_34 | ttc**GAAGAC**tcGCTTgcttggatgatactttaattg | att**GAAGAC**tgagcggctcctctagataccatcaaga |
| pAOX1 genotyping | gactggttccaattgacaagc | |
| pPICZ B genotyping | | gcaaatggcattctgacatcc |
| sfGFP genotyping | | gttcctgcacatagccttcc |
| ManS genotyping | gagcaggagtcaggctcatc | tattgccatcttcctcaccag |
| Venus genotyping | | gcatcaccttcaccttcacc |
| mRuby2 genotyping | gatacgaagatgggggtgtc | caaatgacctccaccatcaa |
| ScGOS1 genotyping | attctaacccggcaatctcc | |
| OCH1 genotyping | cgttttgtacggaccctcac | cgtccagtccatcacatcac |
| pGAP genotyping | tggtttctcctgacccaaag | |
| pFDH1 genotyping | ttcagtgtgctgacctacacg | |
| pDAS1 genotyping | gcctctaggcaaattctgttc | |
| pCAT1 genotyping | atgattttggcctgatgagc | |
| eGFP genotyping | | aagtcgtgctgcttcatgtg |
| ScCYC1tt genotyping | | cttttcggttagagcggatg |
| RPP1Btt genotyping | | ggacaagccgtcactttcac |
| THD3tt genotyping | | ttcaagccaatgctcaaatg |
| pAGT572_Nemo genotyping | ggggtaattaatcagcgaagc | gacaactagggccattgcag |

**Supporting Information**
